# Thera-SAbDab: the Therapeutic Structural Antibody Database

**DOI:** 10.1101/707521

**Authors:** Matthew I. J. Raybould, Claire Marks, Alan P. Lewis, Jiye Shi, Alexander Bujotzek, Bruck Taddese, Charlotte M. Deane

**Affiliations:** Oxford Protein Informatics Group, Department of Statistics, University of Oxford, 24-29 St Giles’, Oxford, OX1 3LB, UK; Data and Computational Sciences, GlaxoSmithKline Research and Development, Gunnels Wood Road, Stevenage, SG1 2NY, UK; Chemistry Department, UCB Pharma, 216 Bath Road, Slough, SL1 3WE, UK; Roche Pharma Research and Early Development, Large Molecule Research, Roche Innovation Center Munich, DE-82377 Penzberg, Germany; Discovery Sciences Department, AstraZeneca, Granta Park, Cambridge, CB21 6GH, UK

**Keywords:** Antibody Therapeutic Structures, Nanobody Therapeutic Structures

## Abstract

The Therapeutic Structural Antibody Database (Thera-SAbDab; *http://opig.stats.ox.ac.uk/webapps/therasabdab*) tracks all antibody- and nanobody-related therapeutics recognised by the World Health Organisation (WHO), and identifies any corresponding structures in the Structural Antibody Database (SAbDab) with near-exact or exact variable domain sequence matches. Thera-SAbDab is synchronised with SAbDab to update weekly, reflecting new Protein Data Bank entries and the availability of new sequence data published by the WHO. Each therapeutic summary page lists structural coverage (with links to the appropriate SAbDab entries), alignments showing where any near-matches deviate in sequence, and accompanying metadata, such as intended target and investigated conditions. Thera-SAbDab can be queried by therapeutic name, by a combination of metadata, or by variable domain sequence - returning all therapeutics that are within a specified sequence identity over a specified region of the query. The sequences of all therapeutics listed in Thera-SAbDab (461 unique molecules, as of 5^th^ August 2019) are downloadable as a single file with accompanying metadata.

## Introduction

Immunotherapeutics derived from B-cell genes are an increasingly successful and significant proportion of the global drugs market, designed to treat a wide range of diseases (1–3).

Whole monoclonal antibody (mAb) therapies dominate the industry - drugs that mimic natural antibodies by containing two identical variable domain structures with a particular specificity (3). The broader class of monoclonal therapies also includes Fragment antigen binding (Fab) regions (a single arm of a whole antibody), single-chain Fv (scFv) regions (a heavy and light chain variable domain connected by an engineered glycine-rich linker), and single-domain variable fragments. These fragments can be expressed in dimeric form to improve avidity, or conjugated with polyethylene glycol (‘pegylated’) for slower clearance (4), with radioisotopes for diagnostic purposes (5), or with radioisotopes or noxious small molecules/peptides for cytotoxicity (6).

Recent developments in protein engineering have resulted in bispecific immunotherapies, where two distinct variable domain binding sites are incorporated into a single protein. As of June 2019, bispecific mAbs, linked Fabs, linked scFvs, and linked single-domain variable fragments have all been assessed in clinical trials (7).

A primary source of information on immunotherapies is the World Health Organisation (WHO), which publishes biennial ‘Proposed’ (8) and ‘Recommended’ (9) International Nonproprietary Name (INN) lists. These INNs serve as globally-recognised generic names by which pharmaceuticals can be identified. To be granted an INN, applicants must include a full amino acid sequence, the closest V and J gene, the IG subclass, and the light chain type (see *https://extranet.who.int/tools/inn_online_application*). This information, coupled with the $12,000 cost of application (as of June 2019), makes INN lists a useful source of therapies that companies intend to carry forward into clinical trials.

Several databases already harvest this information. Two non-commercial antibody-specific resources are the IMGT Monoclonal Antibody Database (IMGT mAb-DB; *http://www.imgt.org/mAb-DB* (10), and WHOINNIG (*http://www.bioinf.org.uk/abs/abybank/whoinnig*). The Therapeutic Antibody Database (TABS; *https://tabs.craic.com*) is antibody-specific and commercial, also scraping patents for therapies. Other databases not specific to antibodies can also capture WHO information, such as ChEMBL (*https://www.ebi.ac.uk/chembl*), Drug-Bank (*https://www.drugbank.ca*), and KEGG DRUG (*https://www.genome.jp/kegg/drug*).

Most databases supply additional metadata for their therapeutic entries, such as clinical trial status, companies involved in development, target specificity, and alternative names. However, while sequence information is available on each therapeutic summary page, it is not possible to query these databases by sequence, nor to bulk-download relevant sets of therapeutic sequences for direct bioinformatic analysis.

Structural knowledge about both the intended target and the therapeutic lead compound is of high importance for rational drug discovery (11, 12). For example, co-crystal complexes reveal where a drug binds to its target (the surface ‘epitope’), and separately-solved structures enable more accurate docking experiments. It can also assist subsequent development and optimisation, as mutants derived from a known structure should be more accurately homology-modeled than those with no close structural partners (13). The Protein Data Bank (14) (PDB) now contains over 150,000 solved structures, and though it is highly biased towards certain protein classes, many diverse targets of pharmacological interest are represented. A significant fraction of these structures contain antibody variable domains, and these are recorded by the Structural Antibody Database (SAbDab (15); 7047 variable domain structures over 3620 PDB entries as of 18^th^ July 2019). Both IMGT mAb-DB and TABS report a set of known therapeutic structures in the PDB, but their reported structural coverage of therapeutic space is low. For example, neither database reports any known structural information for bispecific immunotherapeutics.

To address these deficiencies, we have created the Therapeutic Structural Antibody Database (Thera-SAbDab; *http://opig.stats.ox.ac.uk/webapps/therasabdab*). We harvest sequences as they are released by the WHO, number them with ANARCI (16), and perform a weekly sequence alignment of all therapeutic variable domain sequences to the sequences of known structures stored in SAbDab. Structures with sequence identity matches of 100%, 99%, and 95-98% are recorded and categorised, with alignments on each therapeutic summary page to show precisely where each near-identical structure differs from the therapeutic sequence.

Thera-SAbDab can be queried by INN, by a combination of metadata, such as INN proposal year, clinical trial status, or target, or by sequence (including over a specified region of the sequence). We make available all therapeutic sequences contained within Thera-SAbDab, alongside metadata, to facilitate further research.

## Data Sources

### Sequence Data

Proposed INN lists (8, 9), published by the WHO, are the source of the majority of sequence information in Thera-SAbDab. These are released biannually (one in January/February and another in June/July) and - since list P95 in 2006 - represent a reliable record of variable domain sequences for all antibody- and nanobody-related therapeutics granted a proposed INN. Of the 129 antibody-related therapeutics proposed before 2006, we were able to find sequence information for 47 (36.4%) through the IMGT mAb-DB (*http://www.imgt.org/mAb-DB/*). Although we continue to search, sequences for the remaining 82 may never be released.

All sequences are then numbered by ANARCI (16), which uses Hidden Markov Models to align input sequences to pre-numbered germline sequences. Assigning a numbering allows users to more easily interpret the significance of mutations in near-identical sequence matches. For example, if the mismatch occurs in the extremities of the framework region, it may be judged to have minimal effect on binding site structure.

### Structural Data

Thera-SAbDab compares all numbered therapeutic sequences to the structures in SAbDab (15), which prefilters the PDB (14) for all structures whose sequences align to B-cell germline genes. As all SAbDab structures are also pre-numbered, the comparison of therapeutics to public structural space is efficient. All the existing functionality of SAbDab (e.g. interactive molecular viewers, and numbered structure downloads) is made easily accessible from Thera-SAbDab search results.

### Therapeutic Metadata

Therapeutic metadata comprises a mixture of inherent characteristics and continually-changing status updates.

Certain static properties can be acquired automatically. For example, light chain type is identified through our ANARCI germline alignment (16), while isotype, INN Proposed and Recommended years, and intended target(s) can be harvested directly from the INN lists. Sequence comparison can also be used to identify where different INN names refer to identical variable domains. Other characteristics, such as which companies are involved in therapeutic development, must be manually curated at the time of deposition.

Time-dependent characteristics for new entries are also manually curated after sequence identification, and thereafter every 3 months. We source clinical trial information, developmental status, and investigated condition data from a range of sources including AdisInsight (*https://adisinsight.springer.com*), Clinical-Trials.gov (*https://clinicaltrials.gov*), and DrugBank (*https://www.drugbank.ca*). These websites are updated more regularly, and so are preferable sources for this time-sensitive metadata; we include these fields in Thera-SAbDab to allow for more pharmacologically-relevant searches, as well as to identify all post Phase-I candidates for inclusion in our five updating developability guidelines (17).

## Contents

As of 18^th^ July 2019, Thera-SAbDab is tracking 558 INNs, representing 543 unique therapeutics. Of the 558 INN names, 473 could be mapped to variable domain sequences (87.1%), representing 461 unique therapeutics with sequence data. 436 were monoclonal therapies (three pairs of which share identical variable domains: avelumab & bintrafusp, losatuxizumab & serclutamab, and radretumab & bifikafusp), and 25 were bispecific therapies. Plotting the cumulative sum of these unique therapeutics by year deposited in a WHO ‘Proposed INN’ list shows an exponential increase since the early 2000’s (Figure 1).

**Fig. 1.**
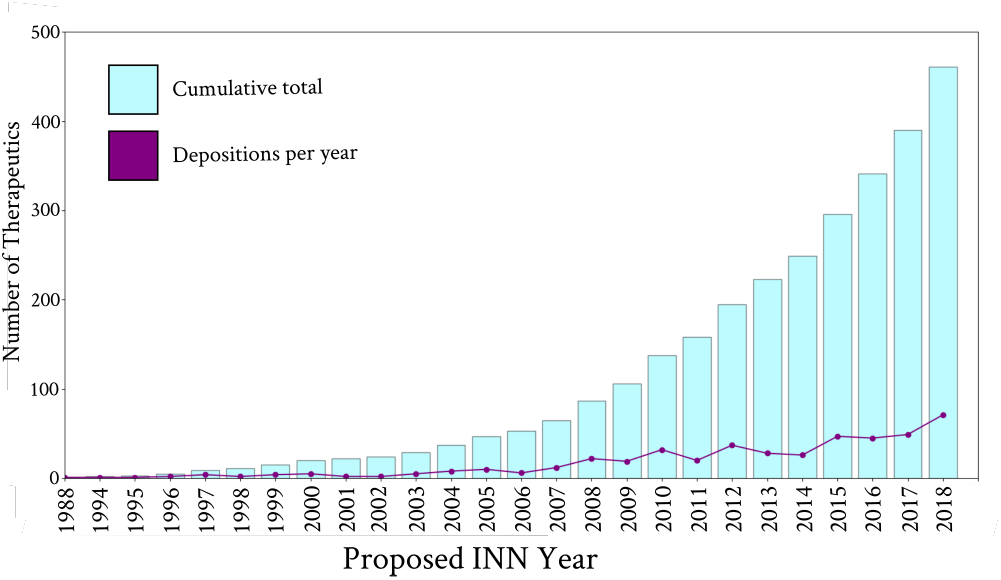
A distribution of the number of antibody- and nanobody-related therapeutics assigned an International Nonproprietary Name (INN) by Year. A record number of 72 of these therapeutics were recognised by the WHO in 2018.

We searched the IMGT mAb-DB (10) and TABS databases (on 28^th^ June 2019) for structures of these 461 therapeutics. IMGT mAb-DB identified 72 structures of therapeutic variable domains, across 36 different monoclonal therapeutics, while TABS reported 53 structures of therapeutic variable domains, across 32 different monoclonal therapeutics. In contrast, Thera-SAbDab (at the 100% sequence identical threshold) recorded 151 therapeutic variable domain structures, across 82 distinct monoclonal therapeutics and 7 distinct bispecific therapeutics. A further 21 monoclonal therapeutics had maximum sequence identity matches of 99% (up to two mutations away from a publicly-available structure), and 12 monoclonals and 4 bispecifics had maximum sequence identity matches of 95-98%. We conclude that, at present, around a quarter (26.4%) of WHO-recognised monoclonal therapeutics have exact or close (> 95% sequence identity) structural coverage. 44.0% of bispecific therapeutics have at least one variable domain with exact or close structural coverage, and two have exact matches for both variable domains.

Thera-SAbDab contains structural information for even the most diversely-formatted therapeutics. Ozoralizumab, a bispecific therapy in active Phase-III clinical trials for rheumatoid arthritis, has a VH(TNFA)-VH(ALB)-VH(TNFA) configuration, where VH(TNFA) is a heavy chain designed to bind to TNF-alpha, and VH(ALB) is another heavy chain designed to bind ALB. Thera-SAbDab has identified a structure for the TNFA binding domain with sequence identity of 95.65% [5m2j; chain D]. Inspection of the sequence alignment shows that 5m2j has a 100% Chothia-defined CDRH3 sequence match to VH(TNFA), and in fact only differs by one mutation across all Chothia-defined (18) CDRs: 31D in VH(TNFA) is 31N in 5m2j. 5m2j is a VHH2 llama nanobody, suggesting that SAbDab’s coverage of nanobody structural space will be increasingly highlighted by Thera-SAbDab as more single-chain therapies arrive in the clinic.

Therapeutically-relevant structures are continually being deposited in the PDB, even many years after initial development. For example, since 2009, the WHO have recorded nine antibody-related therapeutics against IL17A - seven monoclonals and two bispecifics. The first, secukinumab, was recognised in 2009, and since 2014 has been approved for use in certain types of arthritis, psoriasis, and spondylitis. As of early June 2019, there were no close structures for any of these IL17A-binders. However, on 19^th^ June 2019, Eli Lilly deposited an exact variable domain structure for ixekizumab (an IL17A-targetting monoclonal antibody, 6nov) and a close structure for tibulizumab (an IL17A-binding and TNFSF13B-binding bispecific antibody, 6nou) in the PDB (19). SAbDab detected and numbered them in its weekly update, allowing Thera-SAbDab to identify the first publicly-available crystal structure information on IL17A-binding antibodies.

## Usage

There are multiple ways to search Thera-SAbDab. Thera-SAbDab can be queried directly by INN if structural information about a particular therapeutic is needed. Alternatively a combination of metadata can be specified to identify structures for a particular subset of therapeutic space, for example binders to a particular antigen, or therapeutics at a particular stage of clinical trials (Figure 2a). Results are returned in a table format, with links to each therapeutic summary page and a selected array of metadata (Figure 2b).

**Fig. 2.**
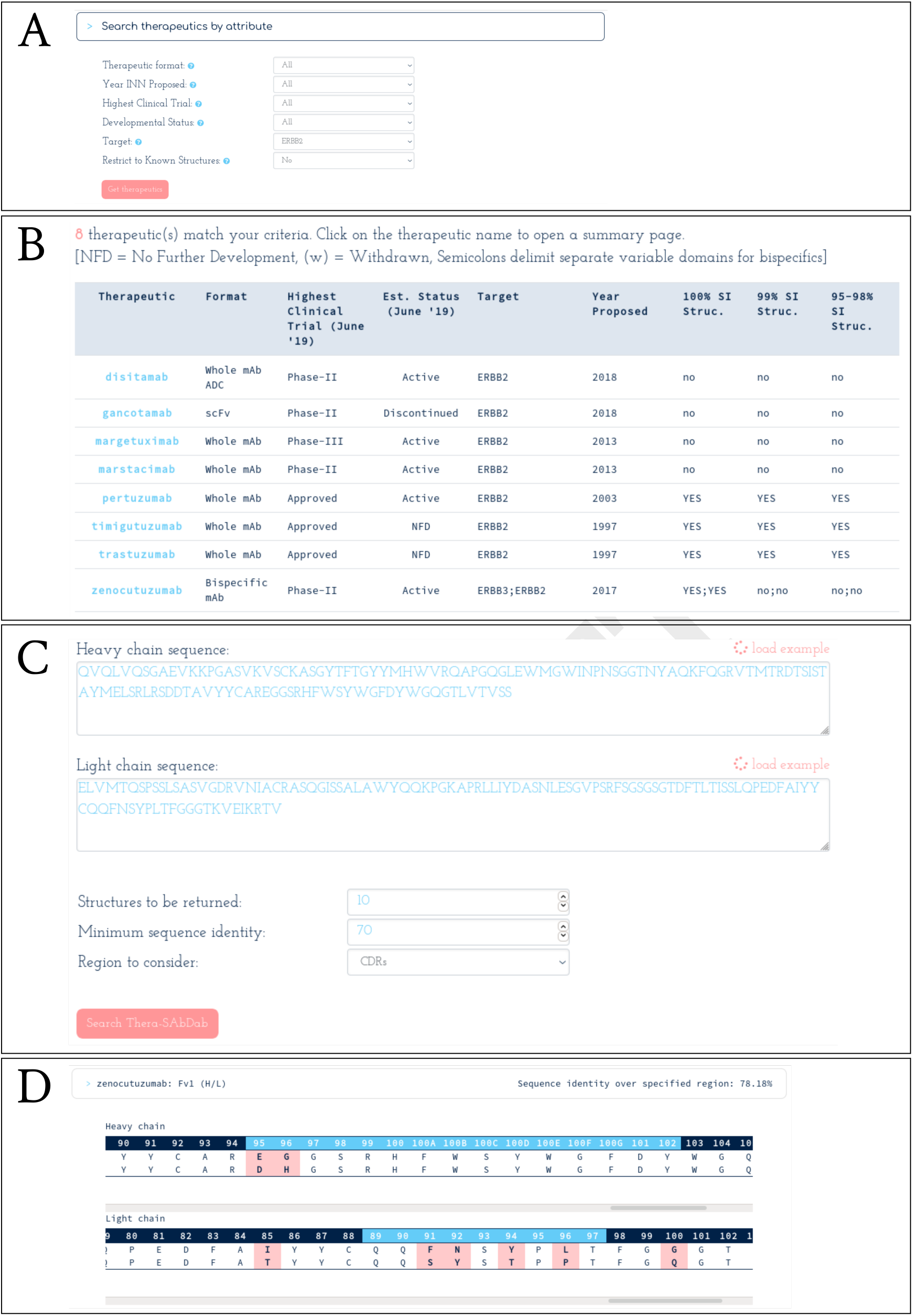
Searching Thera-SAbDab. **(A)** Search by attribute. Here, we search for any therapeutic designed to bind to ERBB2 (often over-expressed in breast cancer). **(B)** Eight therapeutics are designed to bind to ERBB2, seven monoclonals and one bispecific. Four have exact structural information for the ERBB2 binding site. Click the therapeutic name to enter the therapeutic summary page. **(C)** Search by sequence. Here we search for therapeutics with at least 70% sequence identity across the heavy and light chain CDRs of the input sequence. **(D)** Any results are returned alongside sequence identity across the specified region. Alignments show any sequence mismatches across the variable domain sequence.

Each therapeutic summary page lists a structural summary (including our database sequence), with links to relevant SAbDab entries (with PDB codes and chains), and alignment charts (if structures with 95-99% sequence identity are detected). Each SAbDab link redirects the user to the SAbDab summary page for the relevant PDB entry, where all existing functionality can be accessed. Links to appropriate SAbPred

(20) informatics tools (such as ABodyBuilder (21) for variable domain structure modelling, and TAP (17) for developability assessment) are also provided. Finally, we list all the remaining metadata that we have recorded for the therapeutic, ranging from records of investigated conditions, to which companies are developing the therapeutic, to its estimated developmental status.

A third way to search Thera-SAbDab is by sequence (Figures 2c and 2d). This can be harnessed in numerous ways. For example, by querying with a known therapeutic sequence, researchers can look for sequence commonalities between therapeutics over any region of the variable domain. Alternatively, by querying with a developmental candidate sequence, researchers can search for similarity to any other therapeutic, or specifically to those designed to bind to the same target. This could identify potential patenting issues, highlight a risk of polyspecificity, or suggest a binding mode to the intended target.

A further selection of sample use cases for Thera-SAbDab are available at *http://opig.stats.ox.ac.uk/webapps/therasabdab/about*.

## Accessibility

Thera-SAbDab can be queried at *http://opig.stats.ox.ac.uk/webapps/therasabdab*. All sequence data harvested by Thera-SAbDab can be downloaded from the ‘Downloads’ tab of the search page. Sequences are supplied alongside the therapeutic INN, format, isotype, light chain category, highest clinical trial stage reached, and estimated developmental status. We also supply a list of therapeutics for which sequence information has not yet been released.

## Conclusion

We have created Thera-SAbDab with the central aim of collating all public structural knowledge for WHO-recognised antibody- and nanobody-related therapeutic variable domains. Rather than relying on text-mining approaches, which can miss PDB depositions that omit reference to the structure’s therapeutic relevance, Thera-SAbDab uses a systematic approach at the level of sequence identity to detect exact and close matches to our repository of therapeutic variable domains.

This approach has not only enabled us to identify over twice the number of monoclonal therapies with 100% sequence-identical structures in the PDB than in existing databases, but has also identified exact variable domain structures for several bispecific therapies. Our approach can also distinguish between PDB structures with 100%, 99%, and 95-98% sequence identity matches. Sequence alignments guide the interpretation of structures of near-identical sequence.

Like IMGT-DB, Thera-SAbDab can be queried by metadata, but uniquely it can also be queried by variable domain sequence. This enables researchers to identify any therapeutics proximal over any variable domain region to their query sequence.

Thera-SAbDab’s sequence database will be updated with new sequence information twice per year, in line with the release of new WHO Proposed INN lists. An updated list of all therapeutic variable domain sequences with metadata is supplied as a single file to facilitate further analysis.

As shown for IL17A-binding therapeutics, new clinically-relevant structures are continually being released. Accordingly, Thera-SAbDab checks SAbDab after each weekly update for new matches, ensuring that this data is rapidly captured.

## Bibliography

1. A.L. Grilo and A. Mantalaris. The increasingly human and profitable monoclonal antibody market. Trends Biotechnol., 37(1):9–16, 2018. doi: 10.1016/j.tibtech.2018.05.014.

2. S. Steeland, R.E. Vandenbroucke, and C. Libert. Nanobodies as therapeutics: big opportunities for small antibodies. Drug Discov. Today, 21(7):1076–1113, 2017. doi: 10.1016/j.drudis.2016.04.003.

3. H. Kaplon and J.M. Reichert. Antibodies to watch in 2019. mAbs, 11(2):219–238, 2019. doi: 10.1080/19420862.2018.1556465.

4. S. Jevševar, M. Kusterle, and M. Kenig. “PEGylation of Antibody Fragments for Half-Life Extension” in Antibody Methods and Protocols. Methods in Molecular Biology (Methods and Protocols), vol 901. Humana Press, 2012.

5. M. Steiner and D. Neri. Antibody-radionuclide conjugates for cancer therapy: Historical considerations and new trends. Clin. Cancer Res., 17(20):6406–6016, 2011. doi: 10.1158/1078-;0432.CCR-11-0483.

6. A. Beck, L. Goetsch, C. Dumontet, and N. Corvaïa. Strategies and challenges for the next generation of antibody-drug conjugates. Nat. Rev. Drug Disc., 16:315–317, 2017. doi: 10.1038/nrd.2016.268.

7. A.F. Labrijn, M.L. Janmaat, J.M. Reichert, and P.W.H.I Parren. Bispecific antibodies: a mechanistic review of the pipeline. Nat. Rev. Drug Disc., 2019. doi: 10.1038/s41573-019-0028-1.

8. Proposed International Nonproprietary names (INN) List 120. WHO Drug Information, 32 (4), 2018.

9. Recommended International Nonproprietary names (INN) List 81. WHO Drug Information, 33(1), 2019.

10. C. Poiron, Y. Wu, C. Ginestoux, Ehrenmann F., P. Duroux, Lefranc, and M.-P. IMGT/mAb-DB: the IMGT database for therapeutic monoclonal antibodies. JOBIM 2010, 13, 2010.

11. R.L.M. van Montfont and P. Workman. Structure-based drug design: aiming for a perfect fit. Essays Biochem., 61(5):431–437, 2017. doi: 10.1042/EBC20170052.

12. M.I.J. Raybould, W.K. Wong, and C.M. Deane. Antibody-antigen complex modelling in the era of immunoglobulin repertoire sequencing. Mol. Syst. Des. Eng., 2019. doi: 10.1039/C9ME00034H.

13. M.T. Muhammed and E. Aki-Yalcin. Homology modeling in drug discovery: Overview, current applications, and future perspectives. Chem. Biol. Drug Des., 93(1):12–20, 2019. doi: 10.1111/cbdd.13388.

14. H.B. Berman, J. Westbrook, Z. Feng, G. Gilliland, T.N. Bhat, H. Weissig, I.N. Shindyalov, and P.E. Bourne. The protein data bank. Nuc. Acids Res., 28(1):235–242, 2000. doi: 10.1093/nar/28.1.235.

15. J. Dunbar, K. Krawczyk, J. Leem, T. Baker, A. Fuchs, G. Georges, J. Shi, and C.M. Deane. SAbDab: the structural antibody database. Nuc. Acids Res., 42(D1):D1140–D1146, 2014. doi: 10.1093/nar/gkt1043.

16. J. Dunbar and C.M. Deane. ANARCI: antigen receptor numbering and receptor classification. Bioinformatics, 32(2):298–300, 2016. doi: 10.1093/bioinformatics/btv552.

17. M.I.J. Raybould, C. Marks, K. Krawczyk, B. Taddese, J. Nowak, A.P. Lewis, A. Bujotzek, J. Shi, and C.M. Deane. Five computational developability guidelines for therapeutic antibody profiling. Proc. Natl. Acad. Sci. USA, 16(2):4025–30, 2019. doi: 10.1073/pnas.1810576116.

18. B. Al-Lazikani, A.M. Lesk, and C. Chothia. Standard conformations for the canonical structures of immunoglobulins. J. Mol. Biol., 273(4):927–948, 1997. doi: 10.1006/jmbi.1997.1354.

19. R.J. Benschop, C.-K. Chow, Y. Tian, J. Nelson, B. Barmettler, S. Atwell, D. Clawson, Q. Chai, B. Jones, J. Fitchett, S. Torgerson, Y. Ji, H. Bina, N. Hu, M. Ghanem, J. Manetta, V.J. Wroblewski, J. Lu, and B.W. Allan. Development of tibulizumab, a tetravalent bispecific antibody targeting BAFF and IL-17A for the treatment of autoimmune disease. mAbs, 2019. doi: 10.1080/19420862.2019.1624463.

20. J. Dunbar, K. Krawczyk, J. Leem, C. Marks, J. Nowak, C. Regep, G. Georges, S. Kelm, B. Popovic, and C.M. Deane. SAbPred: a structure-based antibody prediction server. Nuc. Acids Res., 44(W1):W474–W478, 2014. doi: 10.1093/nar/gkw361.

21. J. Leem, J. Dunbar, G. Georges, J. Shi, and C.M. Deane. ABodyBuilder: Automated antibody structure prediction with data-driven accuracy estimation. mAbs, 8(7):1259–1268, 2016. doi: 10.1080/19420862.2016.1205773.

